# Unique Trophoblast Chromatin Environment Mediated by the PcG Protein SFMBT2

**DOI:** 10.1101/590356

**Authors:** Priscilla Tang, Kamelia Miri, Susannah Varmuza

**Affiliations:** Department of Cell and Systems Biology, University of Toronto, 25 Harbord St., Toronto, Ontario, Canada, M5S 3G5

## Abstract

Stem/progenitor cells are maintained by a chromatin environment, mediated in part by Polycomb group (PcG) proteins, that depresses differentiation. The trophoblast-specific PcG protein SFMBT2 is known to be required for maintenance of trophoblast progenitors. Rather than binding to trophoblast specific genes repressed in TSC, SFMBT2 is concentrated at chromocenters and regions rich in repetitive elements, specifically LINE sequences and major satellites, suggesting that it is involved in higher order organization of the trophoblast genome. It is also found enriched at a subset of ncRNAs. Comparison of ChIP-seq data sets for other chromatin proteins reveals several stereotypical distribution patterns, suggesting that SFMBT2 interacts with several different types of chromatin complexes specific to the trophoblast lineage.

## INTRODUCTION

Epigenetic biochemistry is as old as cells are old. It has had billions of years to diversify, which is why it is complex, and as evolutionarily varied as it is conserved (Willbanks et al., 2016; Pikaard CS & Mittelsten Scheid O, 2014; Skinner MK, 2011). Our understanding of the role played by epigenetics in life the universe and everything, by comparison, is rudimentary, although the development of new tools has accelerated investigation beyond the blunt hammer of DNA methylation.

Pioneering genetic studies in fruit flies have revealed that gene silencing is rooted in complexes of chromatin proteins (Lewis EB, 1978; Struhl G, 1981; Jürgens G, 1985). One class that has received particular attention is the PcG of proteins, named after one of the first identified gene silencers, Polycomb, required to maintain posterior spatial expression of homeotic genes in fruit fly larvae (Lewis PH, 1949; Lewis EB, 1978; Jürgens G, 1985). The characteristic phenotype – heterozygous ectopic development of sex comb bristles and homozygous lethality due to ectopic expression of homeotic genes – is common to a particular class of proteins which constitute subunits of multi-protein Polycomb Repressive Complexes (PRC1 and PRC2), in addition to other complexes such as PrDUB (Polycomb repressive complex de-ubiquitinase) and PHO-RC (Pleohomeotic repressive complex) (Chittock EC et al., 2017; Schwartz YB & Pirrotta V, 2013; Golbabapour S et al., 2013). In vertebrates, multiple flavours of PRCs have evolved, in part as a result of two whole genome duplication events (Dehal P & Boore JL, 2005; Senthilkumar R & Mishra RK, 2009). This greatly expands the possibilities with respect to developmental innovation.

One such innovation is the placenta. Extraembryonic membranes in egg laying vertebrates evolved into complex structures in mammals that provide an interface between the developing embryo and the mother, a necessity dictated by the exceedingly small mammalian zygote; there isn’t enough mass in a mammalian egg to support development of an entity that can survive and feed. Mammalian development therefore takes place in a specialized extraembryonic environment. While there is growing epidemiological evidence that insults during fetal life can have lasting effects on health (Cluckman PD and Hanson MA, 2004; Langley-Evans SC et al., 1999; Sasaki A et al., 2014; Nakayama M., 2017), there is surprisingly modest interest in examining the role played by the placenta in shaping the fetal developmental environment (Perez-Garcia et al., 2018; Price EM et al., 2016; Zhang X et al., 2015; Suter M et al., 2011; Banister CE et al., 2011).

One area that could benefit from more focused study is the epigenome of the placenta. The ENCODE project at present is greatly enriched by numerous studies of either embryonic stem cells, or neural stem cells (Mouse ENCODE Consortium et al., 2012). Very little work has been done to date on the cells that form the placenta. The study we describe in this paper aims to fill some of that deficit.

Placentas are formed from specialized cells called trophoblast (Rossant J & Cross JC, 2001; Cross JC et al., 2003; Cross JC, 1998). In mouse embryos, trophoblast cells are one of three early lineages to form before the embryo implants in the mother’s uterine wall, the other two lineages represented by embryonic cells and primitive endoderm cells (Cross JC, 1998; Cross JC et al., 1994). In mice, these three types of cells can be induced to form specialized stem or progenitor cells in vitro – trophoblast stem cells (TSC), embryonic stem cells (ESC) and extraembryonic stem cells (XEN) (Hemberger M et al., 2004; Bibel M et al., 2004; Kruithof-de Julio M et al., 2011). One kind of genetic analysis that has helped define the molecular underpinnings of the trophoblast lineage is the ability of mutant embryos to make TSC. Our lab recently reported that the PcG gene *Sfmbt2* is required for the establishment of TSC, indicating that it may form part of the epigenetic framework that supports trophoblast, and by extension development of the placenta (Miri K et al., 2013). *Sfmbt2* was designated a PcG gene because the fruit fly orthologue, *dSfmbt,* displays a classic polycomb phenotype when mutated (Klymenko et al., 2006). Mammalian SFMBT2 can therefore be reasonably expected to form a complex with other chromatin proteins, and to associate with the genome.

ENCODE datasets comprise, largely, whole genome sequencing analyses of the occupation of sites by a variety of chromatin proteins (ChIP-seq) and chromosome conformation capture (Hi-C) (ENCODE Project Consortium et al., 2007; ENCODE Project Consortium et al., 2012; Mouse ENCODE Consortium et al., 2012; Dixon JR et al., 2012). These have allowed investigators to develop novel hypotheses about genome function at a larger scale than had been possible even 10 years ago (e.g. topologically associating domains, or TADs and distal enhancers defined by transcription factor occupancy in specific cell types) (Pope BD et al., 2014; ENCODE Project Consortium et al., 2012; Zacher et al., 2017). Trophoblast cells are just beginning to acquire epigenomic descriptors. In our study, we analyse the occupancy of the genome in TSC by SFMBT2, a protein known from our genetic studies to be required for the maintenance of the stem cell pool. ChIP-seq analyses have revealed that SFMBT2 may play a unique role in the architecture of TSC, mainly through binding to specific repetitive elements, and association with pericentromeric heterochromatin.

### FLAG-tagged TSC

The lentiviral construct expressing myc-tagged SFMBT2 with a linked GFP transgene (Miri et al., 2013) was modified by substitution of the myc-tag with a triple FLAG tag, following digestion of the lentiviral plasmid with BstBI and XbaI, and ligation of a FLAG containing oligonucleotide with engineered sticky ends. Lentiviruses were produced as described (Miri et al., 2013) and used to infect *(C57BL6* X *Castaneus)* F1 TSC at early passage, kindly supplied by Dr. Terry Magnuson.

GFP expressing cells were sorted by fluorescence activated cell sorting (FACS) for three different levels of GFP – strong, medium and weak. The resulting cell pools were expanded into “bulk” cell lines. The experiments described in this paper were generated with the strongly expressing TS cell line designated 7H2-1. Immunohistochemistry revealed that FLAG distribution was indistinguishable from endogenous SFMBT2 (Fig. S1A). Following fixation in neutral buffered 10% formalin (Sigma #HT501320) for 15 min, cells rinsed with phosphate buffered saline (PBS) and permeabilized in 0.3% Triton-X for 15 min. Blocking was done in 10% goat serum and 0.1% Triton-X in PBS for 1 hr at room temperature. Antibodies were diluted in antibody dilution buffer consisting of 5% goat serum and 0.1% Triton-X in PBS. Washes were performed in PBS and 0.05% Tween20. Counter staining was done using 4’,6-diamidino-2-phenylindole (DAPI) at a concentration of 0.1 μg/ml in PBS. Human anti-CREST antibody was purchased from Antibodies Incorporated (#15-234); anti-FLAG antibody was purchased from Sigma-Aldrich (M2, #F3165); anti-SFMBT2 antibody was produced in house (Miri et al, 2013).

### Chromatin immunoprecipitation

#### Harvesting TSC

A minimum of 10^7^ cells is required for a standard ChIP experiment coupled with next-generation sequencing (ChIP-seq). TSC form colonies in culture which minimizes cellular differentiation. However, an accurate cell count requires the dissociation of TSC which may result in stress-induced differentiation or cell death. To circumvent this problem, we calculated the weight of 10^7^ cells, and used this value as a proxy for cell counting. Cells were trypsinized twice with a recovery period of 1 day between trypsin treatments. TSC were washed PBS and incubated with 0.25% trypsin at 37°C for 2 min. Four volumes of standard media were added to neutralize trypsin function and the solution was agitated through pipetting up and down to break cell-cell interactions. Cells were pelleted and resuspended in TS media, plated, and incubated overnight to minimize cellular stress. The trypsin treatment was repeated using a pre-weighed tube and the cell resuspension was used for cell counting by hemocytometer. The remaining cells were pelleted and weighed. 10^7^ TSC weigh approximately 0.23g.

#### Cross-linking with formaldehyde

Plates of TSC were washed 3 times with cold PBS and fixed with 1% formaldehyde at room temperature for 10 min. Formaldehyde was quenched through addition of glycine to a final concentration of 0.125 M at room temperature for 5 min. Cross-linked cells were washed 3 times with cold PBS and harvested in cold PBS supplemented with phenylmethylsulfonyl fluoride (PMSF) and protease inhibitors using a rubber cell scraper. Harvested cells were then aliquoted into pre-weighed tubes for a final weight of 0.23 g. Cells were flash-frozen using a dry ice-ethanol slurry and stored at −80°C until use.

#### Chromatin immunoprecipitation by sonication (SChIP)

Frozen TSC pellets were thawed on ice for 30 mins and chromatin precipitation was performed as previously described by Rada-Iglesias *et al.* (2011). Briefly, chromatin was sonicated to an average size of 75 bp – 400 bp. Sonication parameters were as follows: total processing time of 5 min, amplitude of 40, pulse duration of 10 s, and cooling duration of 30 s. Sonicated chromatin was incubated overnight at 4°C with a 5 μl aliquot of anti-SFMBT2 antibody (Miri et al., 2013), then with 100 μl protein G Dynabeads (ThermoFisher Scientific #10009D) for immunoprecipitation of endogenous SFMBT2 for 2 hrs at 4°C, or with 40 μl FLAG-conjugated beads for immunoprecipitation of FLAG-SFMBT2 (Sigma Aldrich #M8823). Beads were washed 5X with cold RIPA buffer and once with cold TE buffer supplemented with 50 mM NaCl. After phenol-chloroform extraction, aqueous fractions were heated at 55°C for 5 min to remove residual phenol-chloroform then cooled to room temperature for 2 min prior to an ethanol-based DNA precipitation. Ethanol washed DNA pellets were heated at 37°C for 5 min to remove residual ethanol. Samples were then incubated with nuclease-free water at room temperature for 2 min and vortexed. Resuspended DNA was quantified by PicoGreen and stored at −20°C.

#### Chromatin immunoprecipitation by micrococcal nuclease digestion (MN ChIP)

Cells were thawed on ice for 30 mins and chromatin immunoprecipitation was performed as previously described (Tsankov et al., 2015) until the wash procedure. Pellets were resuspended in 1 ml lysis buffer and incubated on ice for 1 hr. Chromatin fragmentation was achieved using 418 units of micrococcal nuclease (MN; Worthington Biochemical Corporation #LS004797) with incubation at 37°C for 2.5 hrs. Digestion was inhibited by addition of EGTA until a final concentration of 50 mM. Washes, elution, and DNA purification steps were performed as in SChIP.

#### Library preparation and high-throughput sequencing

ChIP and total input DNA was sent to The Centre for Applied Genomics (TCAG) for library preparation and subsequent next-generation sequencing. Equal amounts (10 ng) of input and ChIP DNA of sonicated endogenous SFMBT2 samples was supplied for library preparation using the Illumina TruSeq protocol. Sonicated FLAG-SFMBT2 and all MN ChIP samples were prepared using the NEB Ultra DNA library preparation protocol. The NEB Ultra DNA library preparation protocol was used because of low ChIP DNA yields. Although all libraries were subject to paired-end sequencing on the Illumina HiSeq 2500 platform, sonicated endogenous SFMBT2 libraries were sequenced at a different time.

#### Ribonucleic acid sequencing (RNA-seq)

*Sfmbt2-null* embryos were generated by intercrossing heterozygous males and females from the *Sfmbt2* gene trap colony (Miri et al., 2013). Embryos were individually dissected into either embryo (genotyping) or extraembryonic (RNA) portions. Homozygous mutant or wild type RNA extracts from 10-20 embryos were pooled and processed for RNA-seq by TCAG using the Illumina TruSeq protocol and paired end next-generation sequencing on the Ilumina HiSeq 2500 platform. Two biological replicates of each were analyzed.

### Bioinformatics ChlPseq

Illumina fastq data files acquired from sequencing by TCAG necessitated the use of computational programs for data analysis due to the dense information content. All bioinformatics programs were run using an Ubuntu 14.04 OS.

#### Quality assessment and adaptor trimming

The wrapper tool TrimGalore was used to facilitate quality assessment and adaptor trimming (Krueger 2015). FastQC was used for assessing the quality of sequencing reads, identification of adaptor sequences, and identification of non-adaptor overrepresented sequences (Andrews 2010). Adaptor sequences of paired-end reads were trimmed using CutAdapt (Martin 2011). Quality of trimmed reads was then reassessed to ensure removal of adaptor sequences.

#### Genome alignment and peak calling

Adaptor-trimmed sequences were aligned to the mm10 *Mus musculus C57Bl6* genome annotation via Bowtie2 with the quality-check filter implemented (Langmead & Salzberg 2012).

Narrow peaks were called using the MACS2 program with a false discovery rate (FDR) of 0.01 (Zhang et al. 2008). Broad peaks were called using SICER v1.1 with a FDR of 0.01, a window of 200 bp, and a gap of 1000 bp (Zang et al. 2009). Prior to invoking SICER, modifications were made to the GenomeData.py code for inclusion of the mm10 genome. A statistical comparison of genome-wide count distributions was performed using the University of California Santa Cruz (UCSC) utility wigCorrelate (Table S1) (Jee et al. 2011). Due to high correlation between replicates, replicates were pooled and subject to peak calling. All subsequent analyses were performed using SICER-called peak sets.

#### Assessing peak conservation across samples

The R package DiffBind was used for comparison of called peaks across samples (Stark & Brown 2011). DiffBind enabled depiction of similarity between called peaks in a heatmap and generated a consensus peak set. Venn diagrams were generated for visualization of peak conservation in the consensus peak set for each pooled sample and conserved peaks between samples. The fraction of total reads found in called peaks (FRiP) was also calculated.

#### Statistical assessment of SFMBT2 peak association with genomic features

The R package regioneR was used to assess the association between SFMBT2 peak sets with genomic repeats and ncRNA (Gel et al. 2016). Genomic coordinates for mm10 repeats were acquired from RepeatMasker through the UCSC table browser function. Coordinates in the genomic repeat file were binned according their respective repeat families. Information in each repeat family bin was then formatted into BED files for use in regioneR. The same approach was used for ncRNA coordinates acquired from the NONCODE website (Bu et al. 2012). 1000 permutation tests were run per sample with a seed number of 1.

#### Assessing distribution of available ChIP-seq reads across supplied coordinates

Publicly available ChIP-seq data in TSC were downloaded from NCBI GEO. The associated accession numbers are: CTCF (GSM998993), RNA polymerase II (GSM967644), H3K4me1 (GSM1035385), H3K4me2 (GSM967645), H3K4me3 (GSM1035382), H3K27ac (GSM967654), H3K27me3 (GSM967649), H3K36me3 (GSM967646), Total H3 (GSM967647), and Total H2A (GSM1015786, GSM1015787, GSM1015788), H3K9me3 (GSM1035383), H4K20Me1 (GSM967655). All available mm9 BED files were converted to mm10 through the UCSC liftOver utility with automated processing of headers in command-line. NCBI accession numbers with no available BED files were processed from SRA files. SRA files were processed using the sratoolkit to generate fastq files which were subsequently mapped to the mm10 genome using Bowtie2.

SeqMiner was used for read pile-up assessments across specified genomic coordinates (Ye et al. 2014). The shell command for invoking SeqMiner was modified from ‘java – Xmx2000m – jar seqMINER.jar’ to ‘java –Xmx15000m –jar seqMINER.jar’ for increased memory usage. Genomic coordinates of called SFMBT2 peaks or repetitive elements in BED format were supplied as a reference file. Aligned read files for histone or protein distributions of interest were in BED or BAM format. A seed value of 1 was used for all analyses and analysis window was ±5000 bp from the center of coordinates supplied in the reference file.

### Bioinformatics RNAseq

Fastq files acquired from TCAG were analyzed according to the protocol described by Trapnell *et al.* (2012). Reads were aligned to the C57Bl6 mm10 genome using TopHat2 and subsequent analysis was performed using the Tuxedo suite (i.e. Cufflinks, Cuffmerge, Cuffdiff, and the R package CummeRbund). Data was further analyzed using SeqMonk. General Gene Ontology (GO) analysis was performed using the GO analysis tool at Mouse Genome Informatics (MGI) website (www.informatics.jax.org). Significantly differentially expressed genes identified by CuffDiff were cross-analyzed with a list of genes associated with placenta-specific GO terminologies acquired from EMBL-EBI’s QuickGO (www.ebi.ac.uk/QuickGO/).

### Validation ChIP qPCR

Standard qPCR was performed using 20 pg of sonicated FLAG-SFMBT2 template whereas 50 pg of MN-digested FLAG-SFMBT2 template was used. Standards used consisted of serial dilutions of their respective total input fractions. qPCR was performed using the WISENT advanced qPCR mastermix with supergreen lo-rox reagent (WISENT Bioproducts #800-435-UL). Annealing temperatures for all qPCRs was set at 57°C with 40 amplification cycles.

#### Primer Design

Common peaks called across pooled ChIP samples were identified as potential targets. Genomic sequences were then subject to primer design using the Primer3 program. Potential amplicons corresponding to each putative primer set were identified by Primer-BLAST. Primer pairs which generate a single amplicon within 1000 bp were then tested by end-point PCR using input template. Only primers which gave rise to a single distinct amplicon were used for qPCR. Negative targets were designed in the same manner for regions where no fold-enrichment over input was observed.

#### Major satellite qPCR

Primers used for major satellite qPCR were previously published by Martens et al. (2005). qPCR could only be performed on MN-digested FLAG-SFMBT2 samples; 0.5 pg of template was amplified with 1 μM primer.

### GEO Data sets

High throughput sequence data have been deposited in GEO under reference numbers GSE117880, GSE115087 and GSE117879.

## RESULTS

### Loss of SFMBT2 Results in Up-Regulation of Genes

The mammalian Sfmbt2 gene is orthologous with the fruit fly dSfmbt gene, which has been characterized as a Polycomb Group (PcG) gene because the null phenotype of mutants is classic polycomb (Klymenko T et al., 2006; Alfieri C et al., 2013). It is therefore reasonable to assume that the mammalian Sfmbt2 will act like other PcG genes and be involved in transcriptional repression. This question was addressed by performing RNA-seq analysis of Sfmbt2^-/-^ extraembryonic tissues (e7.5 ectoplacental cone and extraembryonic ectoderm). When compared with wild type littermates, the mutant tissues displayed significant up-regulation of 704 genes; in contrast, only 317 genes displayed down-regulation in mutant tissues (Fig. 1). GO analysis suggests most genes affected by knockout of SFMBT2 have roles in developmental processes, cell organization and biogenesis, stress response, cell cycle and proliferation, and signal transduction. Only 20 of 704 genes (0.03%) of all significantly up-regulated genes and 5 of 317 (0.02%) significantly down-regulated genes have GO terminologies associated with placental development. This may in part reflect the underrepresentation of GO data associated with placenta. The stage of embryogenesis chosen, e7.5, represents a stage that precedes the most dramatic phenotypic differences between mutant and wild type embryos, observed at e8.5 (Miri et al., 2013). At e7.5, mutant embryos are generally normal in appearance, although some may have reduced extraembryonic ectoderm. One would expect that loss of a tissue would be reflected in the transcriptome by loss of transcripts from that tissue. It is therefore interesting that so few genes display down-regulation in mutant tissues.

**Figure 1.**
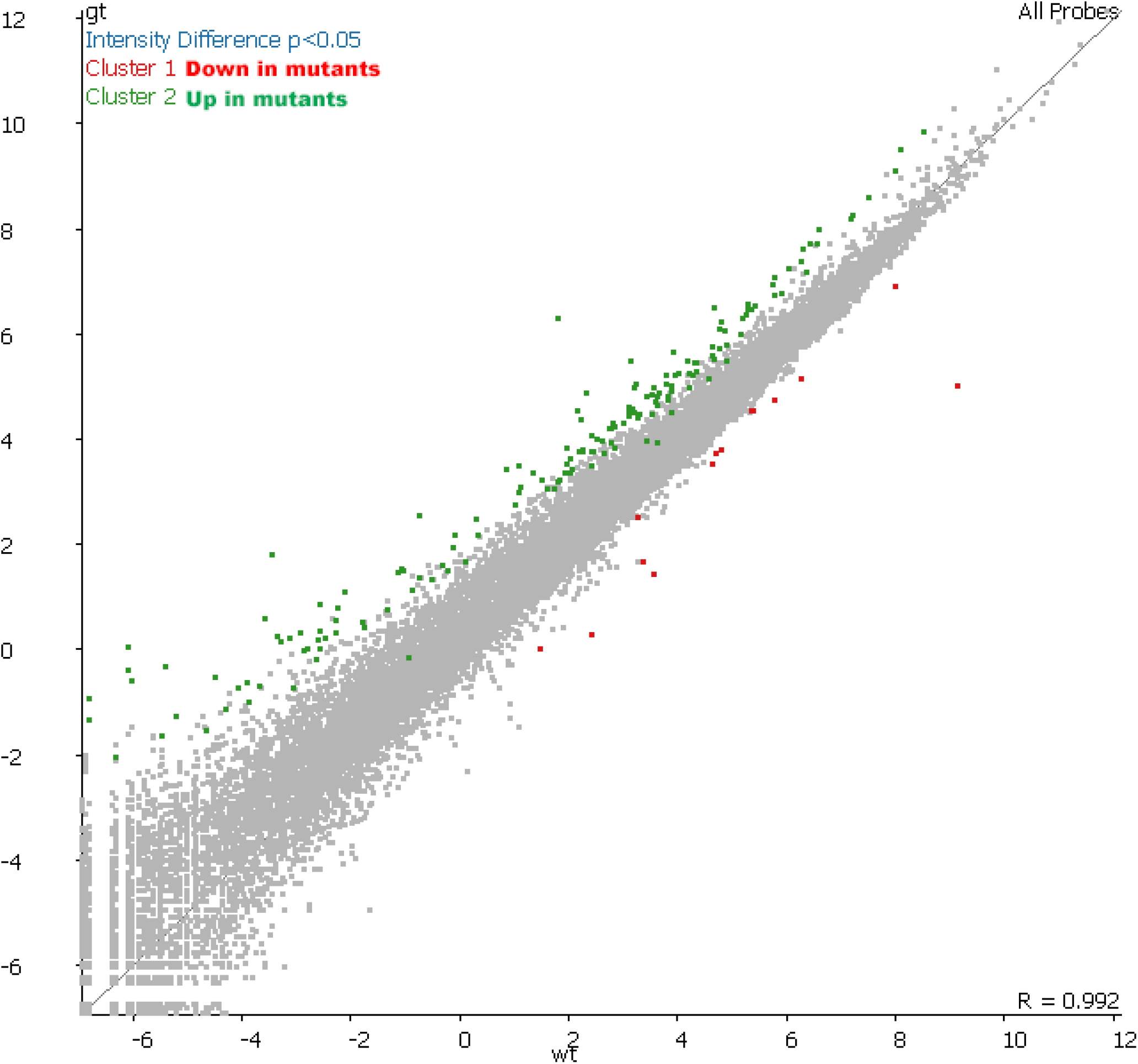
RNA-Seq Analysis of Sfmbt2 Null Extraembryonic Tissues from e7.5 Embryos. Approximately 70% of significantly differentially expressed genes identified in *Sfmbt2-/-* extraembryonic tissues were upregulated (green) compared to wildtype (p<0.05). Few significantly downregulated genes (red) were identified in these tissues. Expression levels of each gene in genetrap mutants (*Sfmbt2gt/gt)* was mapped relative to their wildtype counterparts

### ChIP-Seq Binding of SFMBT2 Displays Broad Peaks

In fruit flies, PcG complexes bind to regulatory sequences called Polycomb Response Elements (PREs) (Kahn TG et al., 2014; Orsi GA et al., 2014; Follmer NE et al., 2012). No clearly definitive PREs have been identified in mammalian cells, suggesting a different mechanism of regulation operates in the larger genomes of vertebrates. In order to assess whether SFMBT2 might be involved directly in transcriptional repression, we performed chromatin immunoprecipitation followed by next generation sequencing in TSC, using both the antibody directed at endogenous SFMBT2 protein and antibody directed against FLAG; the latter was used on cells expressing a FLAG-tagged SFMBT2 transgene.

Initial data analysis with MACS2 did not generate any obvious discrete binding of SFMBT2 to the genome. Given that previous studies showed an interaction between SFMBT2 and modified histones, we subjected the data to analysis with SICER and found that SFMBT2 binds to the genome in a manner similar to that observed for histones, i.e. in modified and broad, peaks. Many of these peaks map to regions rich in repetitive elements, in particular LINE sequences (Fig. 2A). Validation of association with LINE elements was performed by qPCR (Fig. 2B). No obvious peaks were located close to any of the de-repressed genes in mutant extraembryonic tissues.

**Figure 2.**
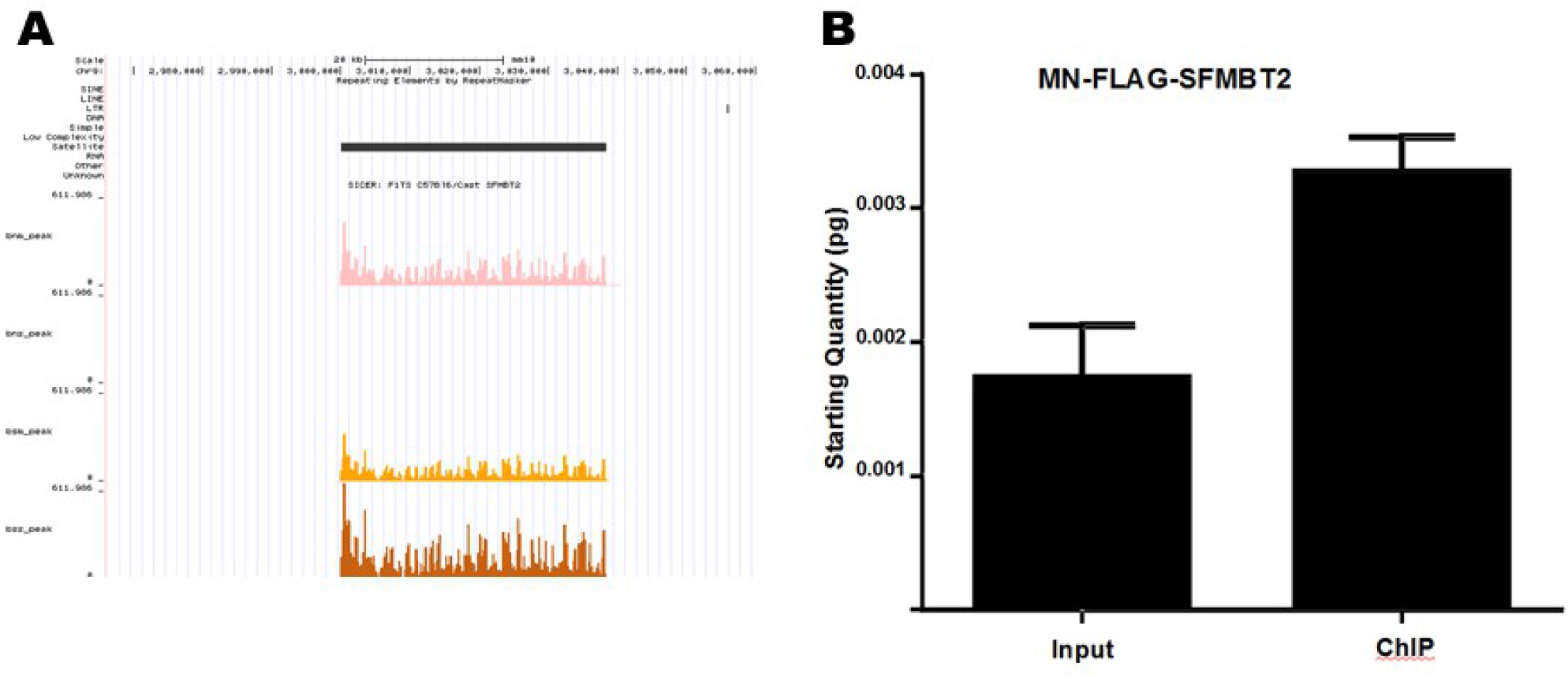
Association of SFMBT2 with Major Satellite Sequences. A. An example of a called SFMBT2 peak localized to region enriched for major satellite sequences. The peak was called using SICER with a FDR of 0.01. Track colours are as follows: MN endogenous SFMBT2 (bnm_peak), pink; MN FLAG-SFMBT2 (bsm_peak), orange; sonicated FLAG-SFMBT2 (bss_peak), brown. Peaks for sonicated endogenous SFMBT2 (bns_peak) were not called for this region likely due to the samples being prepared and sequenced separately. B. Major satellite sequences were significantly enriched in ChIP DNA of MN FLAG-SFMBT2 samples relative to wild-type when equal amounts of DNA were used in qPCR (p<0.005).

### SFMBT2 Associates with Centromeric Major Satellite Sequences and Other Repetitive Elements

Immunohistochemistry in TSC clearly showed an association of SFMBT2 with centromeric regions (Fig. S1B). Because of the repetitive nature of centromeres in mammalian genomes, mapping is problematic. However, during the quality control step in which adapter sequences are removed, we noted that a significant enrichment of major satellite sequences was found in the non-adapter over-represented sequence files in the MN-digested samples of both endogenous and FLAG immunoprecipitates (Table S2). Enrichment was confirmed by qPCR using published primer sequences (Fig. 3).

**Figure 3.**
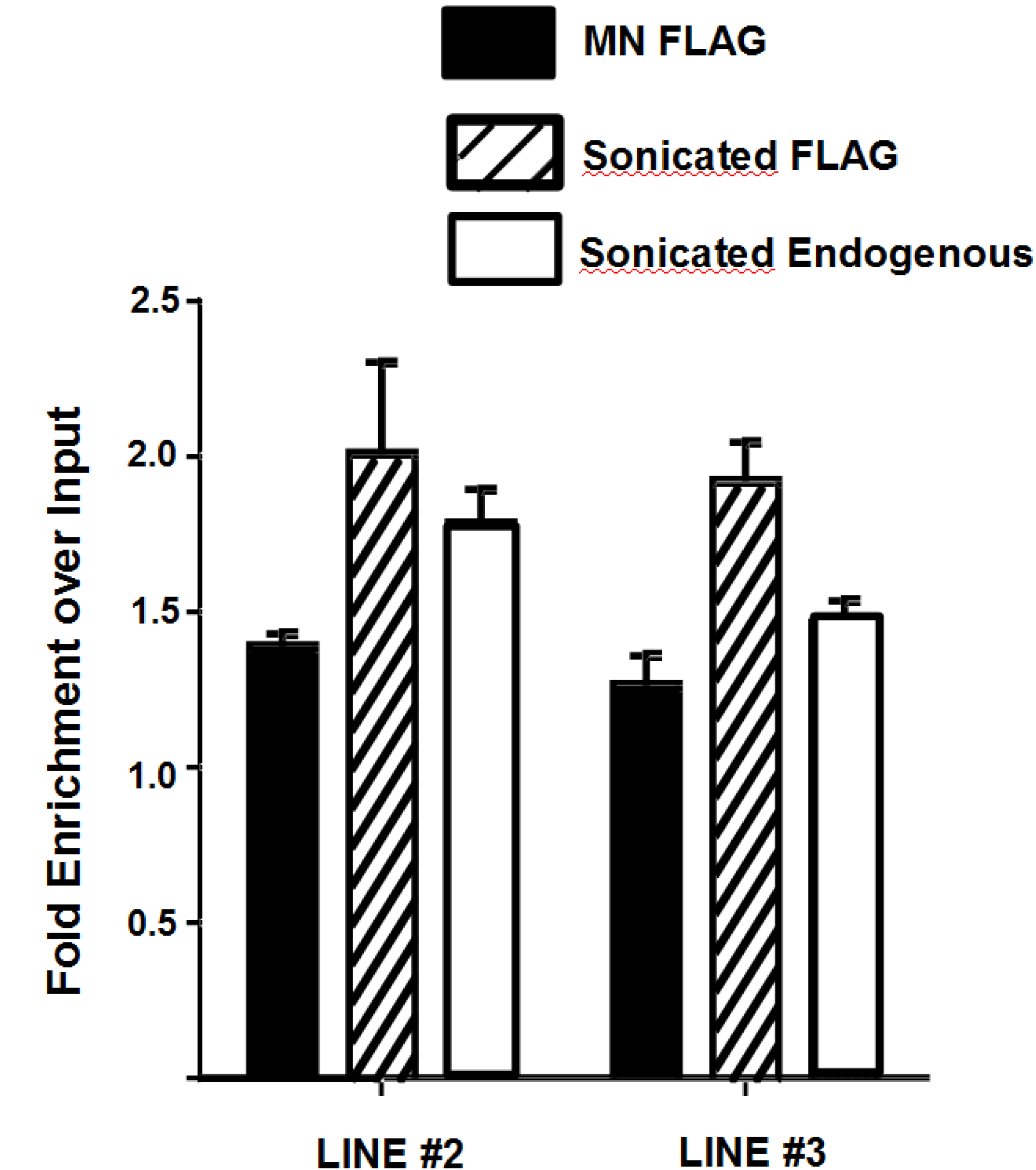
Association of SFMBT2 with LINE Elements. Endogenous and FLAG-SFMBT2 ChIP exhibits enrichment of LINE elements relative to input approximately 1.8- to 2.0-fold even when two different primers were used. Sonicated samples likely display greater fold enrichment due to larger DNA fragments.

Binding of SFMBT2 to LINE elements and major satellite sequences prompted us to look at other classes of repetitive DNA. We used regioneR to visualize the distribution of SFMBT2 in relation to: LINE elements (Fig. 4), low complexity DNA, LTRs, ncRNA, satellite sequences, simple repeats, and SINE elements (Figure S2). Statistically significant patterns were generated for each of these classes of repetitive DNA sequence. Interestingly, association with ncRNAs was quite common (Fig. 5). A statistical analysis of broad peaks using bedtools revealed that between approximately 26% and 46% of SFMBT2 peaks associated with lncRNAs (Table 1), many of which were pseudogenes embedded in regions rich in other repetitive elements, making the design of appropriate qRCR primers problematical.

**Figure 4.**
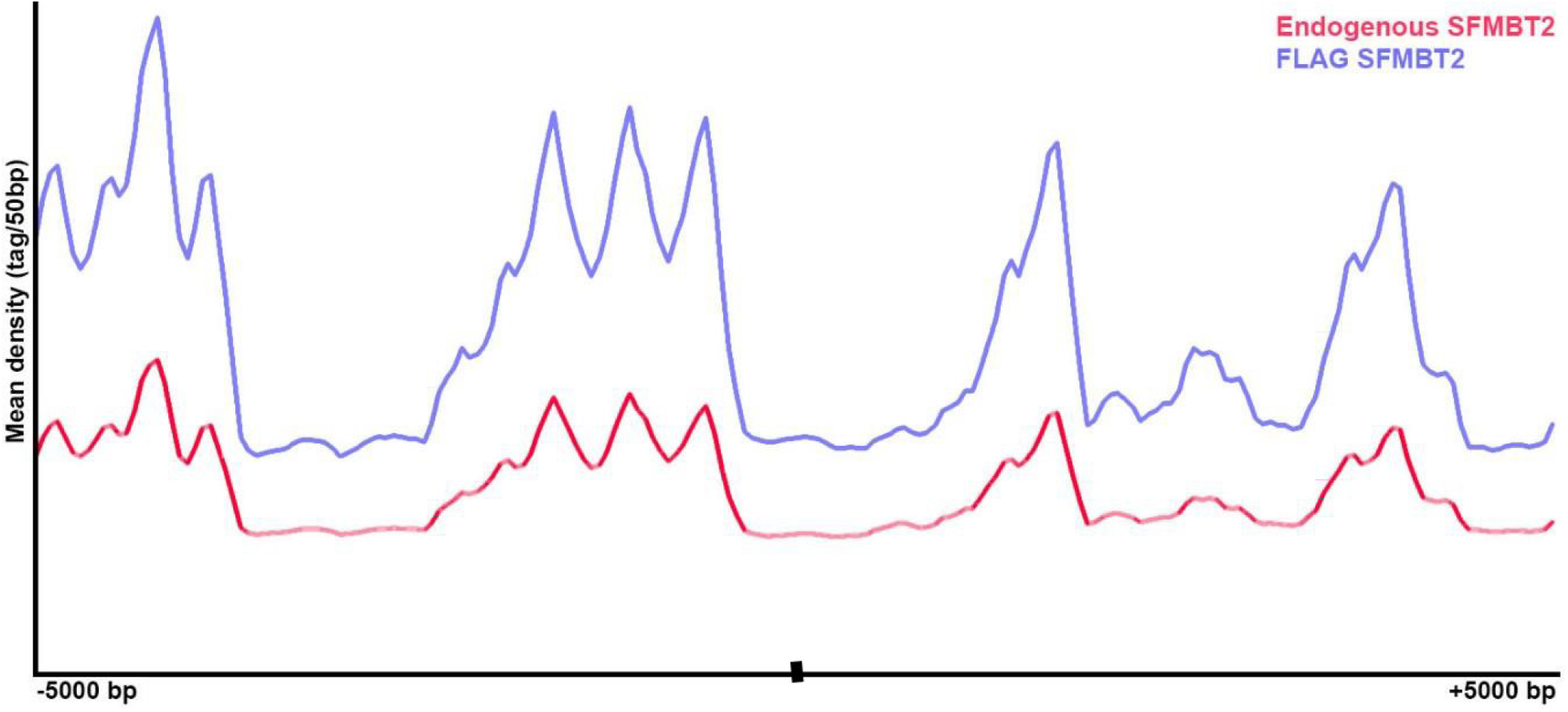
Read Distribution of SFMBT2 Peaks at LINE Elements Across the Genome. RegionR analysis revealed similar patterns of endogenous (red) and FLAG-SFMBT2 (blue) read distributions are observed across known LINE genomic coordinates from 5000 bp upstream to 5000 bp downstream of the center of the specified LINE coordinates.

**Figure 5.**
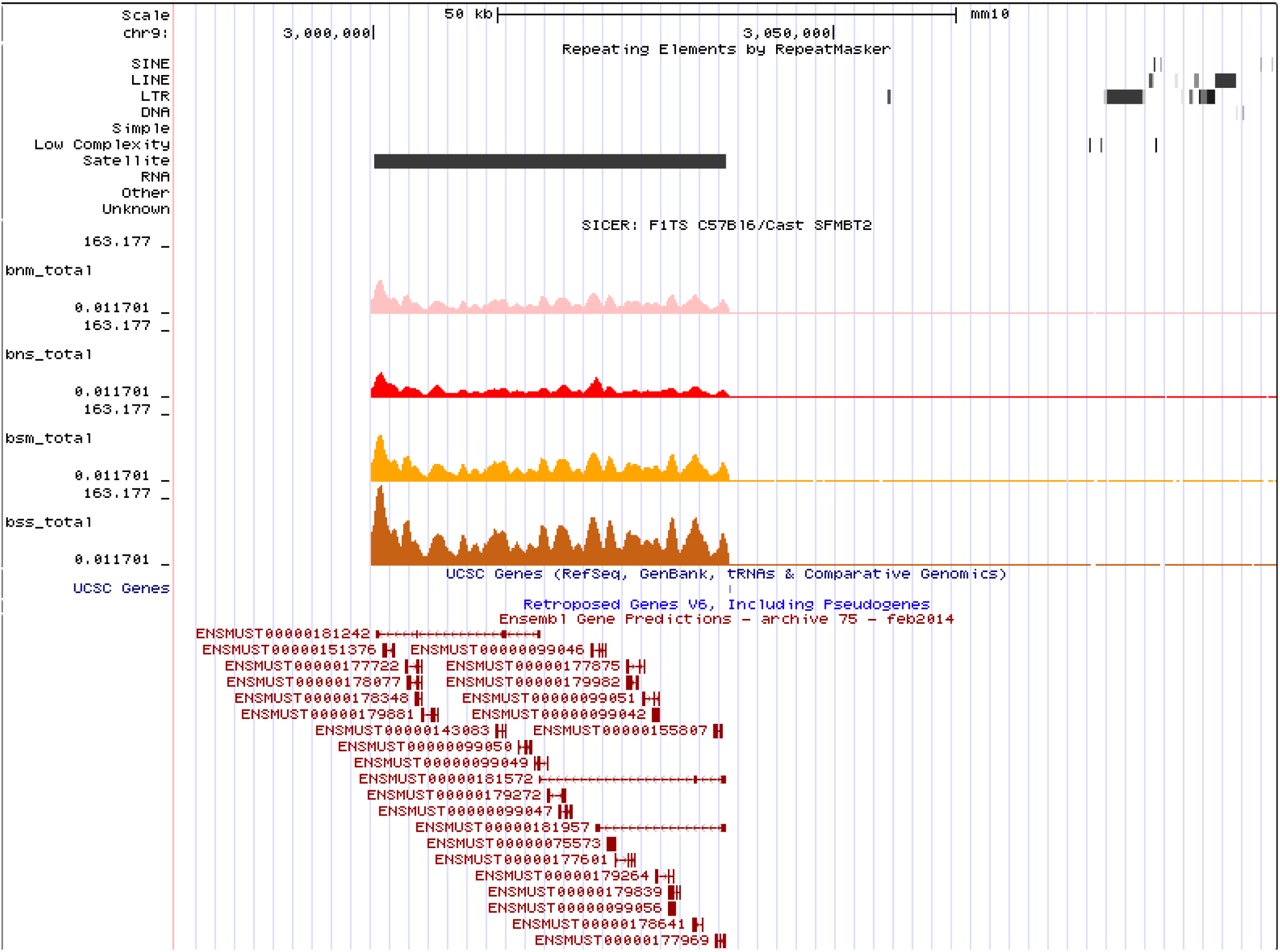
Association of SFMBT2 with ncRNAs. SFMBT2 peaks are mapped to a pericentromeric region enriched for major satellite sequences which also encode for a large cluster of lncRNAs.

**Table 1.** Frequency of overlap between called peaks and known lncRNAs.

| **<u>Sample</u>** | **<u># Peaks</u>** | **<u># lncRNA</u>** | **<u>% Peaks associated with lncRNA</u>** |
| --- | --- | --- | --- |
| MN endogenous | 12350 | 37669 | 46.06 |
| Sonicated endogenous | 2170 | 5673 | 37.50 |
| MN FLAG | 23393 | 63585 | 33.63 |
| Sonicated FLAG | 16087 | 37781 | 26.44 |

#### SFMBT2 Distribution Linked To Stereotypical Histone Marks

The association of SFMBT2 with DNA features such as repetitive and major satellite sequences prompted us to examine the relationship with other histone marks in TSC. Publicly available ChIPseq data for several histone variants were mapped using SEQminer onto the SICER-defined SFMBT2 peaks. Histone H3 displayed a stereotypical distribution surrounding the middle of SFMBT2 peaks, with three strong peaks on one side within about 2.5 kb, and a fourth peak on the other side, at about the same distance (Fig. 6 and S3). The strongest signal is seen in the H3K4me3 data set. A similar pattern is observed in the CTCF data set, while both RNA polymerase II (PolII) and H3K27ac appear to pile up over the SFMBT2 peaks (Fig. 7 and S4). These observations are unexpected, given that H3K4Me3, H3K27Ac and PolII are generally associated with transcriptionally active chromatin, while SFMBT2 appears at least superficially to be transcriptionally repressive (see RNAseq section).

**Figure 6.**
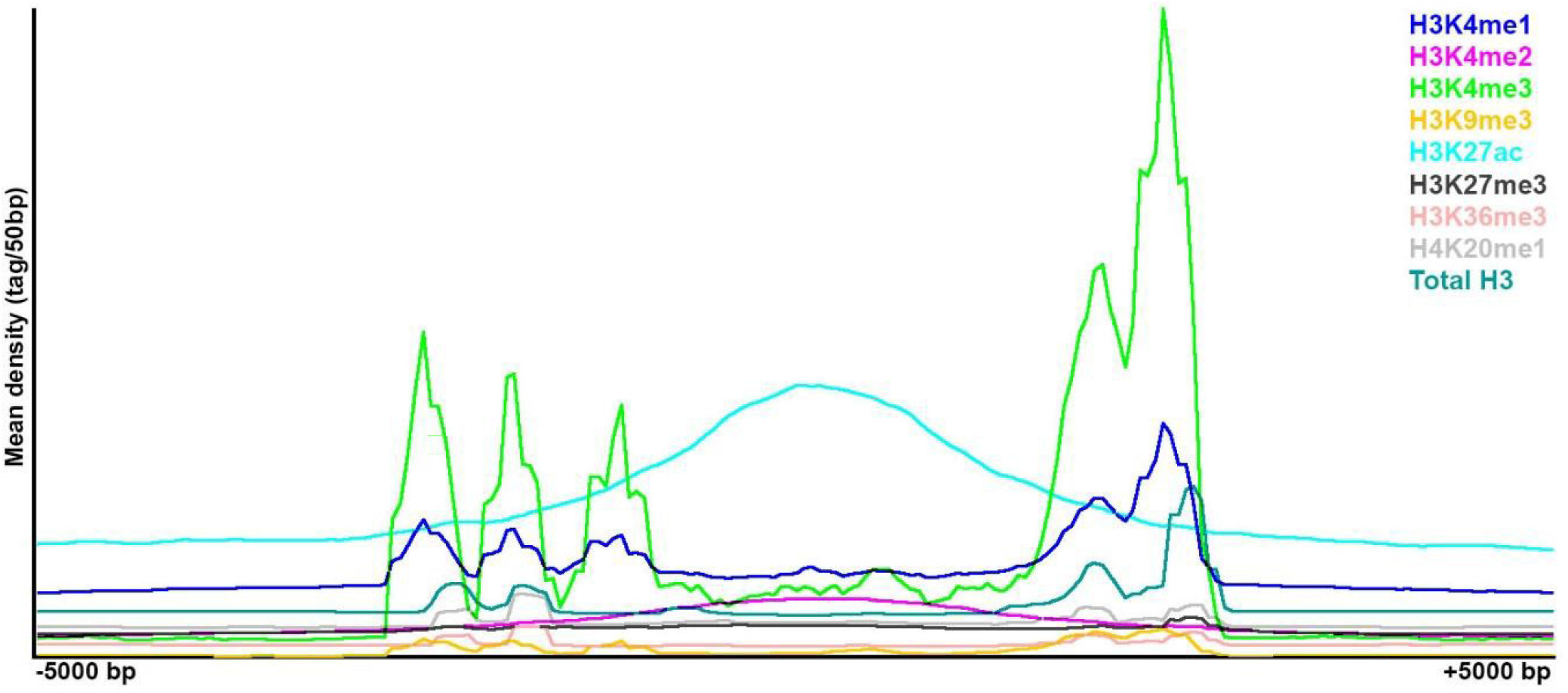
Histone Variant Association with SFMBT2 Peaks. Histones exhibit three distinct patterns of distribution across called endogenous SFMBT2 peak coordinates. The distribution of ChIP-seq reads associated with each histone modification was visualized from 5000 bp upstream to 5000 bp downstream of the center of called SFMBT2 peaks.

**Figure 7.**
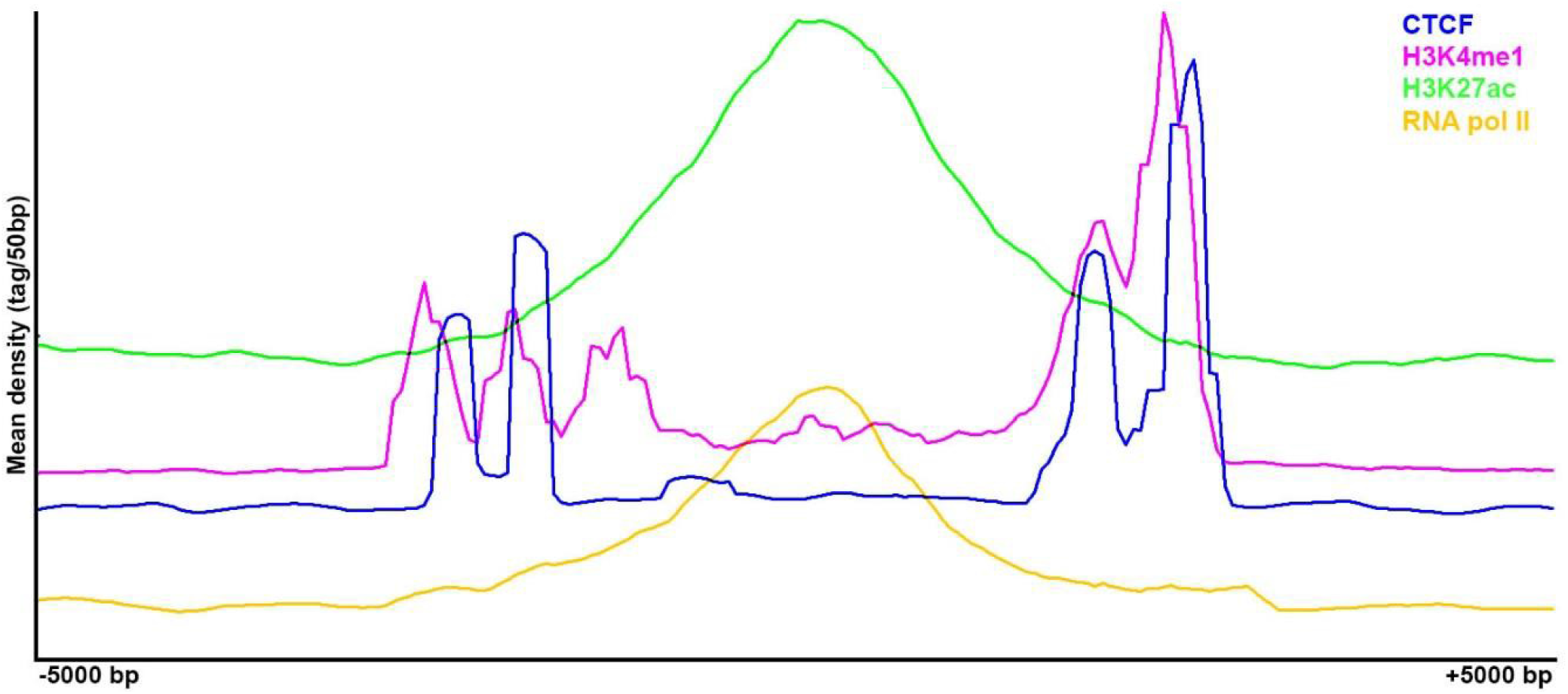
CTCF Association with SFMBT2 Peaks. CTCF and H3K4me1 display different distributions across called endogenous SFMBT2 peak coordinates compared to H3K27ac and RNA pol II. The distribution of ChIP-seq reads associated with each histone modification was visualized from −5000 bp to +5000 bp of the center of called SFMBT2 peaks.

## DISCUSSION

Progenitor cells allow a tissue to bulk up during development or repair itself following injury. Mammals are particularly dependent on stem/progenitor cells during development because embryogenesis is accompanied by an increase in mass, unlike for example insect or amphibian embryos. The variable size of different mammalian species may reflect, at least in part, the renewal capacity of their stem/progenitor cell populations.

The placenta, a highly specialized organ, is dependent, at least in rodents, on the integrity of the trophoblast progenitor population. Reduced numbers of TSC in embryos leads to a reduced placenta and embryonic death. Establishment of TSC has been studied extensively (Ohinata Y & Tsukiyama T, 2014; Rossant J & Cross JC, 2001); however, maintenance of TSC is less well understood. SFMBT2, a PcG protein, is required to maintain the TS compartment. SFMBT2 mutant embryos establish a TS cell compartment, as measured by CDX2 positive cells, but the cell numbers are reduced and placenta growth does not proceed past e8.5 (Miri et al., 2013). Its loss results in de-repression of a suite of genes and premature differentiation of existing TSC into a small placenta. One of the defining features of trophoblast cells is their unusual mode of cell growth following differentiation via a process called endoreduplication, generating several different classes of trophoblast giant cells (TGC) (Simmons et al., 2007). Endoreduplication is characterized by DNA replication in the absence of mitosis, which seemingly makes the requirement for functional centromeres moot. Indeed, we have shown that some TGC in embryos completely lack any SFMBT2 at chromocenters (Miri et al., 2013), and the distribution of SFMBT2 protein in differentiated cells distal to the pool of stem cells at the base of the labyrinth becomes diffuse, further evidence that SFMBT2 function is tied to stemness in trophoblast.

Classic PcG protein complexes in fruit flies regulate target genes by binding to discrete sequences called Polycomb Repressive Elements (PREs), followed by establishment of repressed chromatin. Although a number of genes are de-repressed in SFMBT2 mutant extraembryonic tissues, most are up-regulated by approximately 2-fold, and none has an SFMBT2 peak in the near vicinity; the closest peak is several Mb away. These data support the notion that in mammals, SFMBT2-dependent repression is secondary to the main function, and that the de-repression we observe reflects premature differentiation of extraembryonic tissues. How then is SFMBT2 maintaining the pool of undifferentiated TSC? The distribution of SFMBT2 in TSC is closely associated with features known to be involved in heterochromatin, such as LINE elements, lncRNAs and major satellite sequences. Of note, LINE1 elements have previously been shown to be essential for proper developmental progression and the self-renewal of ESCs in pre-implantation embryos through its role in transcriptional regulation (Percharde M et al., 2018). Perhaps SFMBT2 may be involved in the repression of these LINE elements in TSCs to oppose an ESC phenotype (Nosi U et al., 2017; Roberts RM and Fisher SJ, 2011). Its localization at pericentromeric regions in mitotic cells and chromocenters in interphase cells suggests a strong interaction with heterochromatic elements. At the same time, the stereotypical pile-up of active chromatin marks such as H3K4me3 and H3K27ac to regions flanking SFMBT2 peaks suggests that in TSC, SFMBT2 may be part of the architectural components that maintain the poised chromatin state characteristic of other stem/progenitor cells, and that reduction of SFMBT2 may allow opening up of chromatin during the differentiation process. The stereotypical association of CTCF with SFMBT2 peaks supports this view, given the role this boundary element protein plays in the maintenance of stem cell integrity.

The strongest association, as measured by visualization, is with chromocenters. Our ChIP-seq data contains elements from pericentromeric DNA, e.g. major satellite sequences, and while independent validation using qPCR was successful, it was modest at best, probably a reflection of the technical problems associated with analyzing highly repetitive DNA. Indeed, centromeres are one of the last frontiers in genome analysis (Jain M et al., 2018), their mysteries camouflaged by their repeats. ChIP-seq data sets typically exclude these regions because they are unmappable. This makes pursuit of a structural role in TS cell centromere function highly challenging. Advances in technologies aimed at studying the architectural organization of genomes in undifferentiated, differentiated and abnormal cells has revealed that large distances in the genome are bridged by loops that are defined by Topological Associating Domains (TADs) (Pombo A et al., 2015; Dekker J & Misteli T, 2015; Dixon JR et al., 2012). These features emerge from analysis of multiple chromatin marks in whole genome data sets. Trophoblast stem cells are currently poorly represented in the ENCODE database. The addition of SFMBT2 distribution in trophoblast stem cells reported here will build on a small but growing set of parameters that will allow more comprehensive analysis of this lineage.

## References

1. Alfieri C, Gambetta MC, Matos R, Glatt S, Sehr P, Fraterman S, et al. (2013). Structural basis for targeting the chromatin repressor Sfmbt to Polycomb response elements. Genes & Dev 27:2367–2379

2. Banister CE, Koestler DC, Maccani MA, Padbury JF, Houseman EA, and Marsit CJ. (2011). Infant growth restriction is associated with distinct patterns of DNA methylation in human placentas. Epigenetics 6(7):920–927

3. Bibel M, Richter J, Schrenk K, Tucker KL, Staiger V, Korte M, Goetz M, et al. (2004). Differentiation of mouse embryonic stem cells into a defined neuronal lineage. Nat Neurosci, 7(9):1003–1009

4. Chittock EC, Latwiel S, Miller TCR, and Müller CW. (2017). Molecualr architecture of polycomb repressive complexes. Biochem Soc Trans 45(1):193–205

5. Cross JC, Baczyk D, Dobric N, Hemberger M, Hughes M, Simmons DG, Yamamoto H, et al. (2003). Genes, development and evolution of the placenta. Placenta, 24(2-3):123–130

6. Cross JC, Werb Z, and Fisher SJ. (1994). Implantation and the placenta: key pieces of the development puzzle. Science, 266(5190): 1508–1518

7. Cross JC. (1998). Formation of the placenta and extraembryonic membranes. Ann N Y Acad Sci, 853:23–32

8. Dehal P and Boore JL. (2005). Two rounds of whole genome duplication in the ancestral vertebrate. PloS Biol, 3(10):e314

9. Dekker J and Misteli T. (2015). Long-Range chromatin interactions. Cold Spring Harb Perspect Biol, 7(10):a019356.

10. Deyssenroth MA, Peng S, Hao K, Lambertini L, Marsit CJ, and Chen J. (2017). Whole-transcriptome analysis delineates the human placenta gene network and its associations with fetal growth. BMC Genomics, 18(1):520

11. Dixon JR, Selvaraj S, Yue F, Kim A, Li Y, Shen Y, Hu M et al. (2012). Topological domains in mammalian genomes identified by analysis of chromatin interactions. Nature, 485(7398):376–380.

12. ENCODE Project Consortium, Birney E, Stamatoyannopoulos JA, Dutta A, Guigó R, Gingeras TR, Margulies EH, et al. (2007). Identification and analysis of functional elements in 1% of the human genome by the ENCODE pilot project. Nature, 447(7146):799–816

13. ENCODE Project Consortium, Dunham I, Kundaje A, Aldred SF, Collins PJ, Davis CA, Doyle F, et al. (2012). An integrated encyclopedia of DNA elements in the human genome. Nature, 489(7414): 57–74

14. Follmer NE, Wani AH, and Francis NJ. (2012). A Polycomb group protein is retained at specific sites on chromatin in mitosis. PLoS Genet 8(12):e1003135

15. Gluckman PD, and Hanson MA. (2004). Maternal constraint of fetal growth and its consequences. Semin Fetal Neonatal Med, 9(5):419–425

16. Golbabapour S, Majid NA, Hassandarvish P, Hajrezaie M, Abdulla MA, and Hadi AHA. (2013). Gene silencing and Polycomb group proteins: an overview of their structure, mechanisms and phylogenetics. OMICS 17(6):283–296

17. Hemberger M, Hughes M, and Cross JC. (2004). Trophoblast stem cells differentiation in vitro into invasive trophoblast giant cells. Dev Biol, 271(2):362–371

18. Jürgens G. (1985). A group of genes controlling the spatial expression of the bithorax complex in *Drosophila*. Nature 316:153–155

19. Khan TG, Stenberg P, Pirrotta V, and Schwartz YB. (2014). Combinatorial interactions are required for the efficient recruitment of Pho repressive complex (PhoRC) to Polycomb response elements. PLoS Genet 10(7):e104495

20. Klymenko T, Papp B, Fischle W, Köcher T, Schelder M, Fritsch C, Wild B, et al. (2006). A Polycomb group protein complex with sequence-specific DNA-binding and selective methyl-lysine-binding activities. Genes Dev, 20(9):1110–1122

21. Kruithof-de Julio M, Alvarez MJ, Galli A, Chu J, Price SM, Califano A, and Shen MM. (2011). Regulation of extra-embryonic endoderm stem cell differentiation by Nodal and Cripto signaling. Development, 138(18):3885–3895

22. Langley-Evans SC, Welham SJ, and Jackson AA. (1999). Fetal exposure to a maternal low protein diet impairs nephrogenesis and promotes hypertension in the rat. Life Sci, 64(11):965–974

23. Lewis EB. (1978). A gene complex controlling segmentation in *Drosophila*. Nature 276(5688):565–570

24. Lewis PH. (1949). PC: Polycomb. Drosophila Information Service 21:69

25. Miri K, Latham K, Panning B, Zhong Z, Andersen A, and Varmuza S. (2013). The imprinted polycomb group gene *Sfmbt2* is required for trophoblast maintenance and placenta development. Development, 140(22):4480–4489

26. Mouse ENCODE Consortium, Stamatoyannopoulos JA, Snyder M, Hardison R, Ren B, Gingeras T, Gilbert DM, et al. (2012). An encyclopedia of mouse DNA elements (Mouse ENCODE). Genome Biol, 13(8):418

27. Nakayama M. (2017). Significance of pathological examination of the placenta, with a focus on intrauterine infection and fetal growth restriction. J Obstet Gynaecol Res, doi: 10.1111/jog.13430

28. Ohinata Y, and Tsukiyama T. (2014). Establishment of trophoblast stem cells under defined culture conditions in mice. PLoS One 9(9):e107308

29. Orsi GA, Kasinathn S, Hughes KT, Saminadin-Peter S, Henikoff S, and Ahmad K. (2014). High-resolution mapping defines the cooperative architecture of Polycomb response elements. Genome Res 24(5):809–820

30. Pikaard CS, and Mittelsten Scheid O. (2014). Epigenetic regulation in plants. Cold Spring Hard Perspect Biol, 6(12):a019315.

31. Pombo A, and Dillon N. (2015). Three-dimensional genome architecture: players and mechanisms. Nat Rev Mol Cell Biol, 16(4):245–257.

32. Pope BD, Ryba T, Dileep V, Yue F, Wu W, Denas O, Vera DL, et al. (2014). Topologically associating domains are stable units of replication-timing regulation. Nature, 515(7527):402–405

33. Price EM, Peñaherra MS, Portales-Casamar E, Pavlidis P, Van Allen MI, McFadden DE, and Robinson WP. (2016). Profiling placental and fetal DNA methylation in human neural tube defects. Epigenetics Chromatin 9:6, DOI: 10.1186/s13072-016-0054-8

34. Rossant J, and Cross JC. (2001). Placental development: lessons from mouse mutants. Nat Rev Genet, 2(7):538–548

35. Sasaki A, deVega W, Sivanathan S, St-Cyr S, and McGowan PO. (2014). Maternal high-fat diet alters anxiety behaviour and glucocorticoid signaling in adolescent offspring. Neuroscience, 272:92–101

36. Schwartz YB, and Pirrotta V. (2013). A new world of Polycombs: unexpected partnerships and emerging functions. Nat Rev Genet 14:853–864

37. Senthilkumar R and Mishra RK. (2009). Novel motifs distinguish multiple homologues of *Polycomb* in vertebrates: expansion and diversification of the epigenetic toolkit. BMC Genomics 10:549

38. Skinner MK. (2011). Role of epigenetics in developmental biology and transgenerational inheritance. Birth Defects Res C Embryo Today, 93(1):51–55.

39. Struhl G. (1981). A gene product required for correct initiation of segmental determination in *Drosophila*. Nature 293:36–41

40. Suter M, Ma J, Harris A, Patterson L, Brown KA, Shope C, Showalter Lori et al. (2011) Maternal tobacco use modestly alters correlated epigenome-wide placental DNA methylation and gene expression. Epigenetics 6(11):1284–1294

41. Willbanks A, Leary M, Greenshields M, Tyminski C, Heerboth S, Lapinska K, Haskins K et al. (2016). The evolution of epigenetics: From prokaryotes to humans and its biological consequences. Genet Epigenet, 8:25–36

42. Zacher B, Michel M, Schwalb B, Cramer P, Tresch A, and Gagneur J. (2017). Accurate promoter and enhancer identification in 127 ENCODE and roadmap epigenomics cell types and cells by GenoSTAN. PLoS One, 12(1): e0169249

43. Zhang X, Pei L, Li R, Zhang W, Yang H, Li Y, Guo Y et al. (2015). Spina bifida in fetus is associated with an altered pattern of DNA methylation. J Hum Genet 60(10):605–611

